# Digital Spatial Profiling of Collapsing Glomerulopathy

**DOI:** 10.1101/2021.09.08.459502

**Authors:** Kelly D. Smith, Kammi Henriksen, Roberto F. Nicosia, Charles E. Alpers, Shreeram Akilesh

## Abstract

Collapsing glomerulopathy is a histologically distinct variant of focal and segmental glomerulosclerosis that presents with heavy proteinuria and portends a poor prognosis. Collapsing glomerulopathy can be triggered by viral infections such as HIV and SARS-CoV-2. Transcriptional profiling of collapsing glomerulopathy lesions is difficult since only a few glomeruli may exhibit this histology within a kidney biopsy and the mechanisms driving this heterogeneity are unknown. Therefore, we used recently developed digital spatial profiling (DSP) technology which permits quantification of mRNA at the level of individual glomeruli. Using DSP, we profiled 1,852 transcripts in glomeruli from HIV and SARS-CoV-2 infected patients with biopsy confirmed collapsing glomerulopathy. The increased resolution of DSP uncovered heterogeneity in glomerular transcriptional profiles that were missed in early laser capture microdissection studies of pooled glomeruli. Focused validation using immunohistochemistry and RNA *in situ* hybridization showed good concordance with DSP results. Therefore, DSP represents a powerful method to dissect transcriptional programs of pathologically discernible kidney lesions.

## INTRODUCTION

Collapsing glomerulopathy is a distinct histologic pattern of focal and segmental glomerulosclerosis (FSGS).^1-3^ It is characterized by collapse of the capillary tuft, hypertrophy/proliferation of the overlying epithelial cells, and usually extensive effacement of podocyte foot processes. Collapsing glomerulopathy is considered the most aggressive form of FSGS and is linked to progressive parenchymal injury leading to end stage kidney disease.^4^ Collapsing glomerulopathy is known to be associated with drugs of abuse, bisphosphonates, high interferon states (hepatitis C treatment, lupus) and viral infections (parvovirus B19, HIV, SARS-CoV-2). Individuals with high risk APOL1 genotypes are particularly susceptible to develop collapsing glomerulopathy reflecting an interplay between genetic susceptibility and exogenous triggers.^5-7^

It has been difficult to study molecular mechanisms driving this distinct glomerulopathy given its rarity and the focal nature of collapsing lesions within a kidney biopsy. A seminal laser capture microdissection and microarray-based transcriptomic study demonstrated activation of developmental programs in FSGS and idiopathic collapsing glomerulopathy.^8^ A more recent laser capture microdissection and proteomic study identified changes in extracellular matrix (ECM) components in idiopathic collapsing glomerulopathy.^9^ Both studies required pooling of histologically normal and collapsing glomeruli from multiple sections in order to obtain enough material for the assays. Pooling precludes comparing gene expression between individual glomeruli, interferes with the ability to assign changes to affected vs. unremarkable glomeruli, and masks glomerular heterogeneity.

Due to demanding sample input requirements, acquiring individual glomerular transcriptional profiles in collapsing glomerulopathy biopsies has not been achievable using laser capture microdissection approaches. However, technological advances in spatial transcriptomics now permit accurate measurement of gene expression and localization to defined anatomic structures. One of these technologies, NanoString’s GeoMx digital spatial profiling (DSP) platform, is compatible with formalin-fixed paraffin embedded (FFPE) tissue material and permits free-form selection of anatomic structures. The required minimum capture area of 8-10,000µm^2^ from a single 4µm thick section (30-50 cells) is well suited to profile glomeruli that are typically 200µm in diameter with ≥ 100 cells/glomerulus cross section.

Because it is still not known if a universal mechanism drives virus mediated collapsing glomerulopathy compared to other etiologies, we explored molecular mechanisms driving similar morphologic presentations in SARS-CoV-2 and HIV associated collapsing glomerulopathy using digital spatial profiling of gene expression. This study of gene expression in individual glomeruli provides novel insight into the pathogenesis of collapsing glomerulopathy, and the molecular heterogeneity that exists even among morphologically indistinguishable glomeruli.

## METHODS

### Slide preparation for spatial profiling

This study was approved by the University of Washington and the University of Chicago’s institutional review boards. Three patients with COVID-19 and collapsing glomerulopathy were identified in the course of a previous study.^5^ Three patients with HIV associated collapsing glomerulopathy and three patients with hematuria and preserved kidney function (“normal” controls) were identified via database searches. For spatial transcriptomic profiling, tissue sections from multiple patients were placed on the same slide (3 slides total) to reduce number of slides needed for probe hybridization. 4µm sections of formalin fixed paraffin embedded kidney biopsy tissue were affixed to charged slides and baked at 60°C for 30 minutes. After deparaffinization and rehydration, antigen retrieval was performed by treating slides with boiling Tris EDTA, pH 9 for 15 minutes followed by 1µg/ml proteinase K in PBS for 15 minutes at 37°C. After washing in PBS, Cancer Transcriptome Atlas (1825 targets) and COVID-19 spike in probes (27 targets) were hybridized to the slides overnight at 37°C. Probes directed against each target gene consist of oligonucleotides complementary to multiple portions of the gene’s mRNA which improves target detection in FFPE tissue sections. Each target probe also incorporates a unique photocleavable oligonucleotide barcode that can be quantified by next generation sequencing. The next day, following stringency washes and blocking, fluorescently labeled antibodies to detect CD45 (Novus, 2B11+PD7/26, #NBP2-34528AF647), smooth muscle actin (SMA; Abcam, 1A4, ab202368), pan-cytokeratin (nanoString, AE1/AE3, #121301310) and a DNA binding dye (ThermoFisher, Syto13, S7575) were applied to the slides for 1 hour at room temperature.

### Spatial profiling, sequencing and data analysis

Following washes, prepared slides (without coverslips) were immediately loaded into the nanoString GeoMx Digital Spatial Profiler and covered with acquisition buffer. Glomerular regions of interest (ROI) were selected and outlined by two kidney pathologists (KDS, SA) using the instrument’s web interface. Oligonucleotide barcodes encoding their target gene were released from their target-specific portions using ultraviolet illumination, captured and then sequencing libraries prepared per the manufacturer’s protocol. An average of 916,867 raw reads/ROI (range 303,623-2,393,264) were collected and matched to target genes using nanoString’s custom oligonucleotide barcode-to-gene matching pipeline. Imputed raw read counts per gene were third quartile (Q3) normalized. Parametric ANOVA with p< 0.05 was used as a test for significance without multiple testing correction in order to maximize sensitivity of detection. After the spatial profiling experiment was completed, the slides were stained with Jones methenamine silver for brightfield visualization of tissue sections and correlation of histology with gene expression.

### In situ hybridization and immunohistochemistry

RNA *in situ* hybridization (ISH) for VEGFA (ACDBio #423161) and SARS-CoV-2 (ACDBio #848561) was performed using a RNAscope 2.5 HD kit (ACDBio #322300) with DAB colorimetric development followed by hematoxylin counterstaining. Immunohistochemistry for PAX8 (Sigma MRQ-50) and MME/CD10 (Thermo Fisher 56C6) was performed on a Leica Bond autostainer per protocols established in the clinical laboratory at the University of Washington Medical Center, Seattle, Washington, USA.

## RESULTS

We identified three patients each with HIV-associated and SARS-CoV-2 associated collapsing glomerulopathy. Glomeruli with collapsing features appeared histologically similar across both groups of patients by light microscopy (**Figure 1A**). One COVID-19 patient (COVID3) had prominent thrombotic microangiopathy (TMA) with several glomeruli containing thrombi and others displaying ischemic changes (**Figure 1A**). Three patients with hematuria and preserved kidney function who were found to have normal histology on kidney biopsy were selected as controls. Patient demographics/clinical characteristics are shown in **Supplemental Table 1** and biopsy findings are summarized in **Supplemental Table 2**. Sections were hybridized with Cancer Transcriptome Atlas probes (1,827 genes + 27 spike ins to detect SARS-CoV-2 viral sequences and host genes important for viral entry and reproduction) and immunofluorescence counterstained with markers to detect nuclei, smooth muscle actin (SMA), pan-cytokeratin and leukocytes (CD45) (**Figure 1B**). Individual glomeruli were selected in freeform regions of interest (ROI) and correlated to their histology on Jones-stained sections (**Supplemental Figure 1**). Unfortunately, due to tissue folding a glomerulus with collapsing features was not available for patient COVID3. For each glomerular ROI, oligonucleotide barcodes linked to the bound probes were released using UV illumination and collected for high throughput sequencing. ROI area and sequencing parameters are shown in **Supplemental Table 3**. After 3^rd^ quartile (Q3) normalization, expression profiles were generated at the individual glomerulus level for each patient (**Figure 1C, Supplemental Table 4**). Principal component analysis demonstrated variation by disease and histology with tight clustering of expression signatures from all glomeruli derived from the 3 normal controls (**Figure 1D**). The glomeruli from patient COVID3 with TMA behaved distinctly and were considered separately in all subsequent analyses. These results demonstrated feasibility of spatially registered transcriptomic profiling of individual glomeruli with correlation to histology.

**Figure 1.**
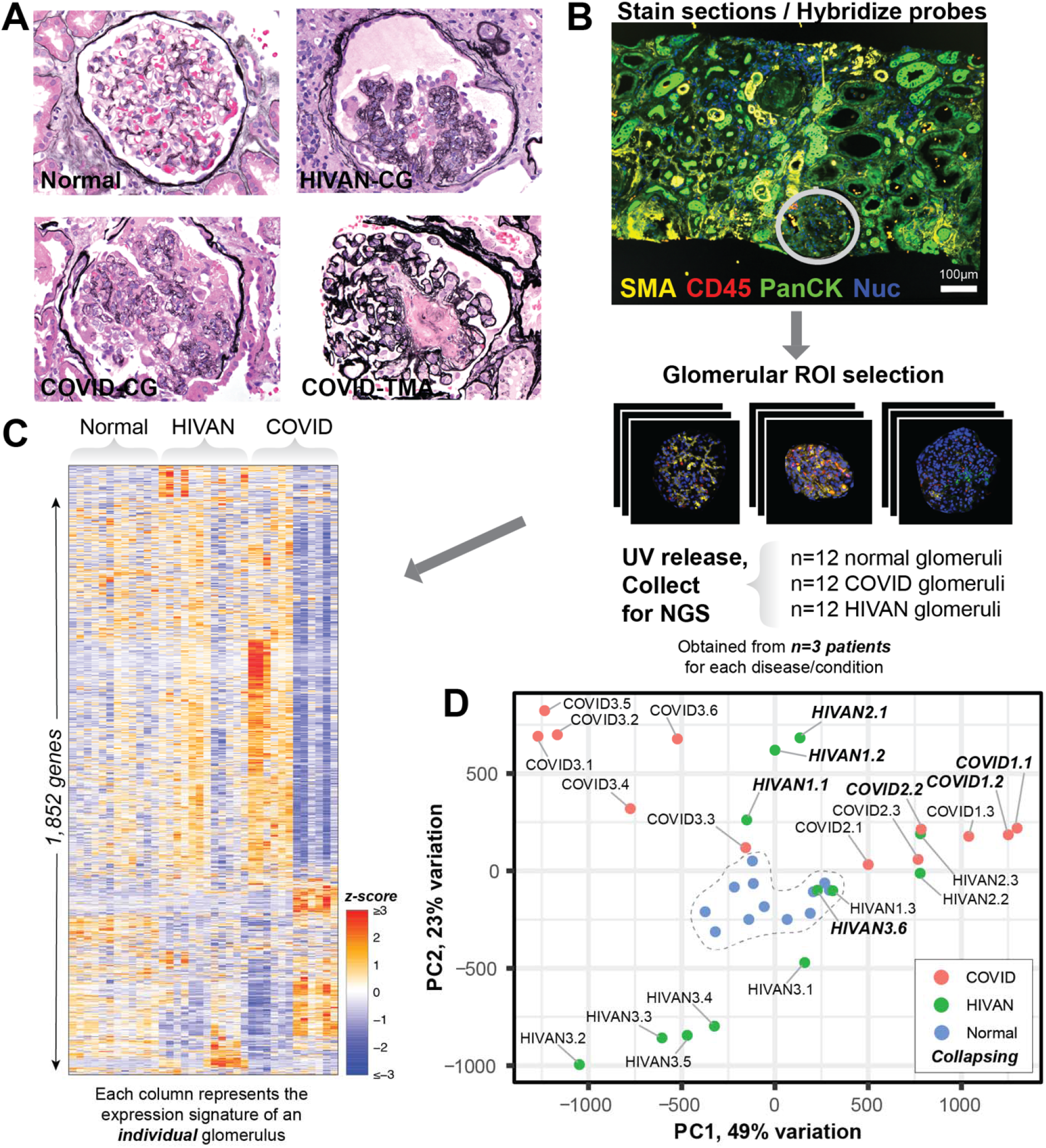
Digital spatial profiling of individual glomeruli from HIV and COVID-19 patients. **A**) Representative Jones-stained sections of glomeruli with histology analyzed by digital spatial profiling. HIVAN – HIV associated nephropathy; CG – collapsing glomerulopathy; COVID – SARS-CoV-2 associated disease; TMA – thrombotic microangiopathy. **B**) Biopsy sections were immunostained with fluorescent antibodies against the indicated markers and free from regions of interest (ROI) were drawn around glomeruli of interest. Bound probes were released and quantified by next generation sequencing. **C**) 1852 genes (rows) were quantified from each glomerulus (columns) from n=3 patients each from normal, HIVAN and COVID conditions. **D**) Principal component analysis reveals tight clustering of normal glomerular transcriptome profiles from 3 patients but large differences amongst glomeruli from HIVAN and COVID patients. For each glomerulus the first number indicates the patient and the second, the glomerular ID for that patient. Glomeruli with features of collapsing glomerulopathy are indicated in ***bold italics***.

Next, we asked how DSP data compared to laser capture microdissection data that required pooling of glomeruli. Leveraging the individual glomerulus-level DSP data, we first asked what gene expression differences characterized glomeruli with collapsing histology regardless of inciting etiology. We compared the expression profiles of 7 glomeruli with collapsing features to 12 glomeruli from the 3 normal controls. This identified 209 genes with >1.5 fold change in expression level with parametric ANOVA p< 0.05 (**Figure 2A**). These differentially expressed genes were enriched in kidney development and morphogenesis ontologies (**Figure 2B**). These enriched ontologies correlated well with a previous laser capture microdissection and microarray-based study of collapsing glomerulopathy^8^ and there was good overlap of differentially expressed genes between the two studies (**Figure 2C**). Shared differentially expressed genes such as *CD24, CDH2* and *PAX8* were increased in collapsing glomeruli (**Figure 2D**). CD24, a progenitor cell marker has been validated in human collapsing glomerulopathy.^10^ CD44^11^ and PAX8^12^ are increased in animal models of FSGS but have not been demonstrated in human collapsing glomerulopathy. Interestingly, many genes that were differentially expressed in our study were decreased in glomeruli with collapsing morphology and have not been reported previously. We found that expression of podocyte genes such as *WT1, CDKN1C* and *MME* (CD10) were reduced in DSP data, but were not significantly changed in the prior laser capture microdissection study,^8^ perhaps due to dilution of signal resulting from pooling glomeruli with collapsing and normal morphology. Genes involved in endothelial homeostasis (*ERG, VEGF, KDR, FLT1*) were also decreased in both types of collapsing glomerulopathy. Mitotic markers *MKI67* and *PCNA* were both significantly increased in collapsing glomeruli in our study, but with < 1.5 fold change (not shown). The loss of podocyte phenotypic markers and gain of proliferative markers is consistent with prior reports of a dysregulated podocyte phenotype in collapsing glomerulopathy.^13-19^ To localize expression changes to specific cell types within the glomerulus, we validated expression and localization of PAX8 and MME in additional sections from our samples. In normal kidney samples and glomeruli from patient COVID3 with TMA, PAX8 showed strong nuclear expression in parietal epithelial cells lining Bowman’s capsule as well as in tubular epithelial cells (both excluded from our ROI selection for DSP) but not in cells on the glomerular tuft.^20^ In contrast, collapsing glomeruli demonstrated strong PAX8 expression in most of the reactive epithelial cells overlying the collapsed capillary tuft as well as sporadic expression in individual podocytes and in parietal epithelial cells lining Bowman’s capsule (**Figure 2E**). MME showed the reverse pattern with reduced expression in collapsing glomeruli compared to patient COVID3 with TMA and the normal controls. This could represent either decreased expression in podocytes or loss/trans-differentiation of podocytes themselves. Therefore, individual glomerular level profiles uncovered transcriptional heterogeneity and novel differentially expressed genes in collapsing glomeruli, which were then validated with protein detection by immunohistochemistry.

**Figure 2.**
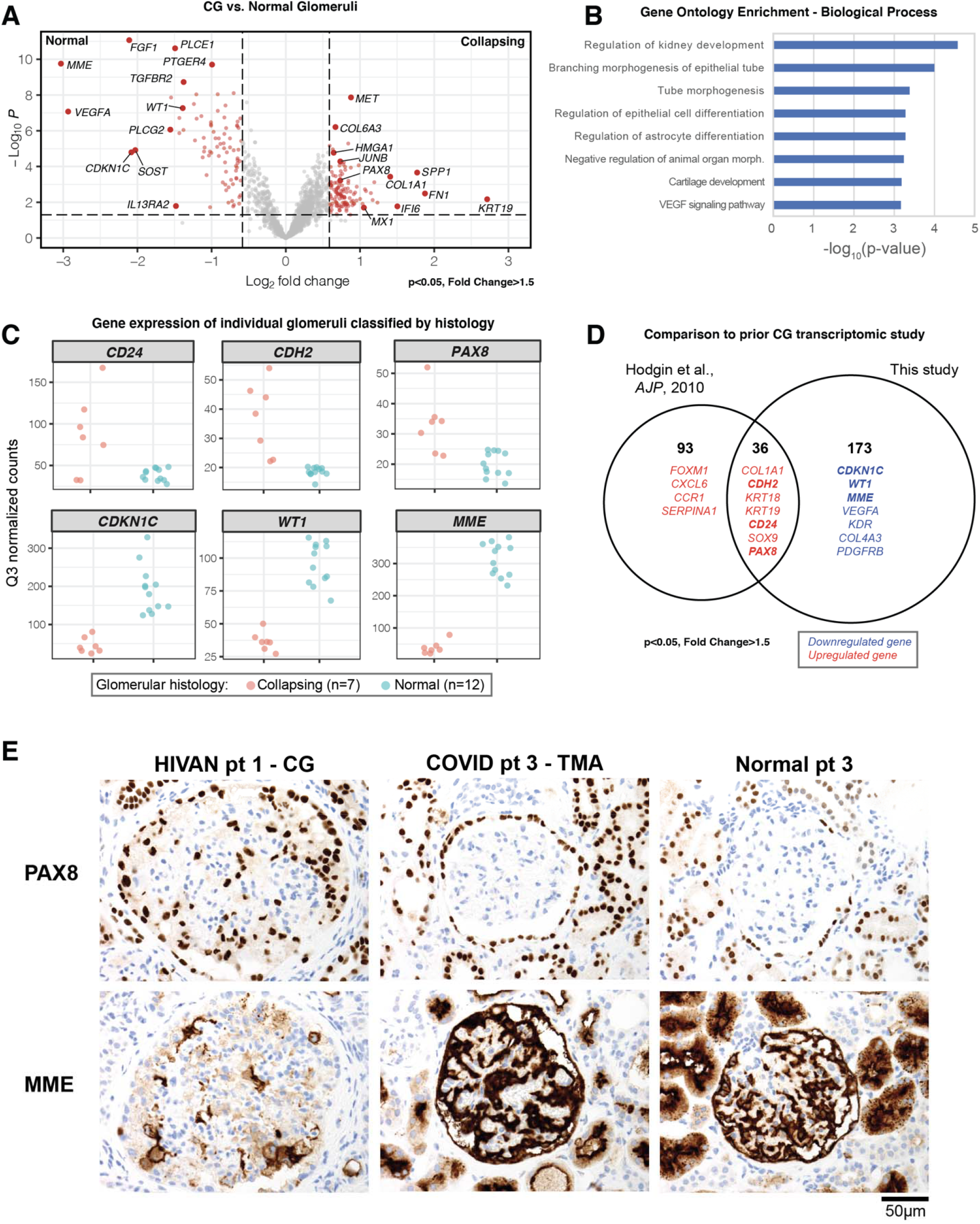
A transcriptomic signature of collapsing glomerulopathy. **A**) Volcano plot showing genes with differential expression between glomeruli with collapsing features and glomeruli from normal patient controls. Parametric ANOVA p< 0.05 and absolute fold change of >1.5-fold were used to identify genes of interest. **B**) Gene ontology enrichment identifies significant changes in biological processes related to kidney development, morphogenesis and VEGF signaling in collapsing glomerulopathy. **C**) Dotplots of gene expression for selected genes. Each dot represents an individual glomerulus classified according to histology. For all comparisons between collapsing and normal glomeruli, parametric ANOVA p< 0.05. **D**) Comparison of our results to a previous transcriptomic study of collapsing glomerulopathy identifies several shared gene signatures. Differences between the studies likely reflect different sensitivities/dynamic ranges of the techniques or dilutional effects in the prior study which pooled all glomeruli from patients with a diagnosis of collapsing glomerulopathy whether they had collapsing features or not. Expression of genes highlighted in boldface are shown in panel C. **E**) Immunohistochemistry for PAX8 and MME validates the glomerular expression profiles of collapsing glomerulopathy shown in panel C.

To understand whether HIV and SARS-CoV-2 affected glomeruli in similar or different ways, we compared the expression profiles of the 12 glomeruli from 3 HIV-infected patients to the 6 glomeruli from patients COVID1 and COVID2 (again excluding patient COVID3 with TMA). This identified 58 genes with >1.5-fold change in expression level with parametric ANOVA p< 0.05 (**Figure 3A**). Of note, SARS-CoV-2 viral RNAs were not detected in any of the glomeruli examined and ISH for SAR-CoV-2 spike protein was negative in all 3 COVID-19 biopsies (*data not shown*), a finding consistent with a recent DSP study of autopsy kidneys.^21^ Gene ontologies related to anti-viral responses and interferon pathways were strongly enriched in glomeruli from patients with HIV (**Figure 3B**). Interestingly, when broken down by histology, it was apparent that both collapsing and histologically normal appearing glomeruli from HIV-infected patients manifested a strong interferon response signature (**Figure 3C**). Our previous analysis had identified *KRT19* as a gene that was upregulated in collapsing glomeruli (**Figure 2A**). However, examination of COVID and HIV collapsing glomeruli revealed that this was driven only by collapsing glomeruli from HIV-infected patients. These increased *KRT19* expression levels correlated well with pan-cytokeratin immunofluorescence performed as a counterstain during ROI selection (**Figure 3D**). Thus even though biopsy findings were morphologically similar in HIVAN and COVID-19 patients, DSP revealed molecular pathways that were differentially regulated in HIVAN and COVID-19.

**Figure 3.**
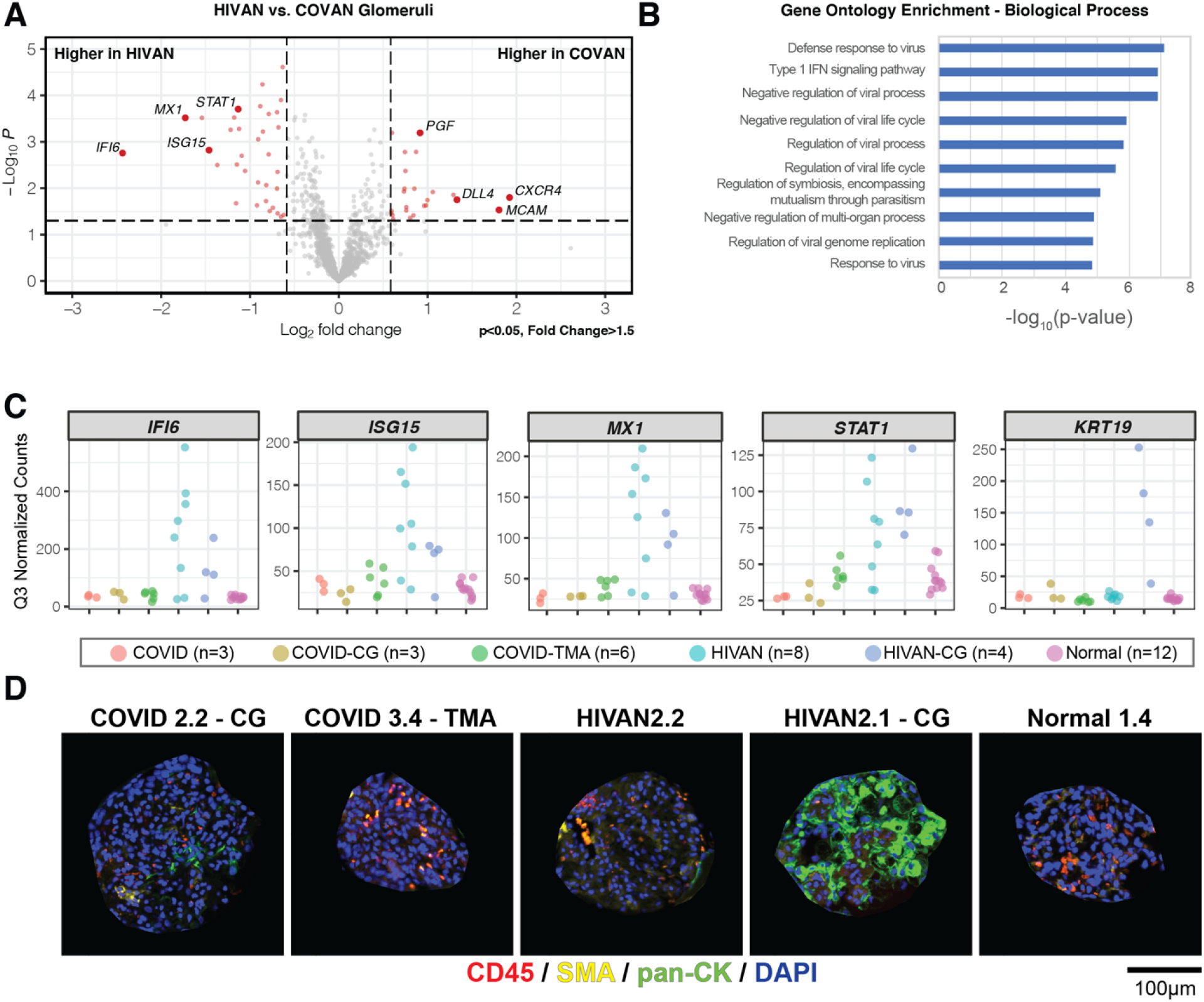
Interferon signaling is a dominant feature of HIVAN glomeruli. **A**) Volcano plot showing genes with differential expression between glomeruli from patients with HIVAN or COVID-19 (n=3 patients each). Parametric ANOVA p< 0.05 and absolute fold change of >1.5-fold were used to identify genes of interest. **B**) Gene ontology enrichment identifies significant changes in biological processes related to anti-viral responses and interferon signaling in glomeruli from HIVAN patients. **C**) Dotplots of gene expression for selected genes. Each dot represents an individual glomerulus classified according to histology. For all comparisons except *KRT19*, parametric ANOVA p< 0.05. **D**) Immunofluorescence images of captured glomerular regions of interest confirms elevated expression of keratins in HIVAN-CG correlating with panel C.

Collapsing glomerulopathy can also be detected in the context of vascular endothelial injury (thrombotic microangiopathy, TMA) such as in patients undergoing treatment with VEGFA blockade for cancer.^22, 23^ TMA and collapsing glomerulopathy have also been reported in association with COVID-19 kidney injury as was the case for patient COVID3 but the underlying mechanisms are poorly understood.^5, 24^ Therefore, we studied the expression profiles of the 6 glomeruli from patient COVID3 with TMA in comparison to the 12 normal glomeruli. In this analysis, 182 genes had >1.5-fold change in expression level with parametric ANOVA p< 0.05 (**Figure 4A**). These genes were enriched for ontologies related to metabolism, transmembrane transport, tissue remodeling and angiogenesis (**Figure 4B**). Strikingly, several genes important for vascular function were increased in glomeruli from patient COVID3 with TMA (**Figure 4C**). These included adhesion molecules VE-cadherin (*CDH5*)^25^ and *PECAM1* (CD31),^26^ the angiopoietin signaling receptor *TIE1*,^27^ Notch signaling components *DLL4* and *NOTCH1*,^28^ and the early stress responsive AP-1 transcription factor component *JUNB*.^29^ Consistent with ischemic injury related to TMA, hypoxia response genes *DDIT4*,^30^ *HIF1A*^*31*^ and *VEGFA*^*32*^ were also induced. The expression patterns of VEGFA classified by glomerular histology were particularly striking. To investigate this further, we performed RNA in situ hybridization (RNA-ISH) for *VEGFA* in additional tissue sections from our patients’ biopsies (**Figure 4D**). Normal glomeruli showed robust podocyte staining for *VEGFA* mRNA. By contrast, collapsing glomeruli from an HIV patient showed almost complete loss of VEGFA staining. Unfortunately, collapsing glomeruli were not present in the deeper levels obtained from the COVID patient biopsies. *VEGFA* levels also appeared lower in non-collapsing glomeruli from both HIV and COVID patients. In contrast, glomeruli from patient COVID3 with TMA exhibited intense podocyte staining for *VEGFA*. These RNA-ISH expression patterns corroborated individual glomerular expression patterns derived by spatial profiling and identified a complex podocyte/endothelial related expression program that varied as a function of glomerular histology.

**Figure 4.**
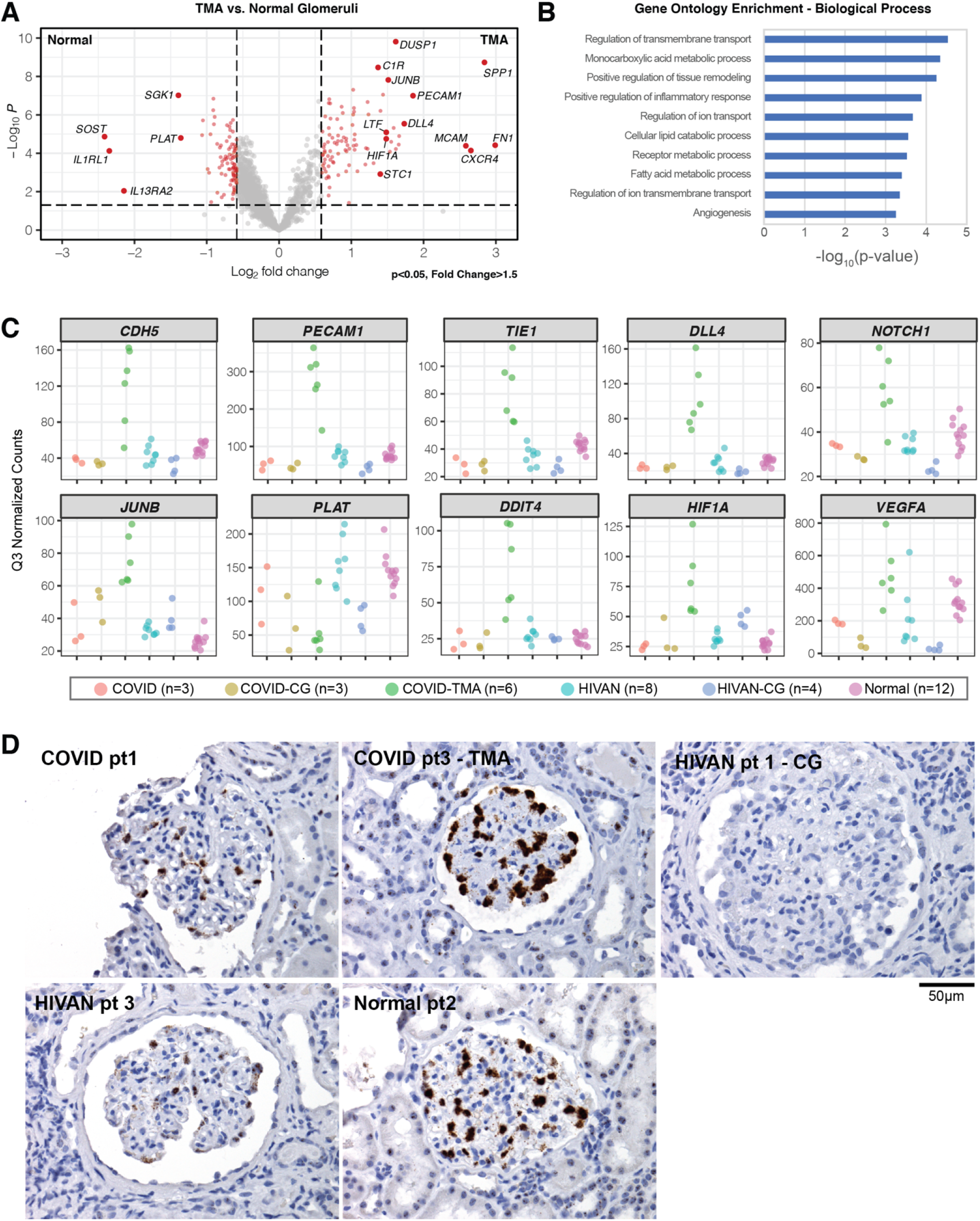
TMA induces a vascular stress response signature in COVID. **A**) Volcano plot showing genes with differential expression between glomeruli from the patient with COVID-TMA (patient 3, n=6 glomeruli) compared to 3 normal patients’ glomeruli (n=12 glomeruli). Parametric ANOVA p< 0.05 and absolute fold change of >1.5-fold were used to identify genes of interest. **B**) Gene ontology enrichment identifies significant changes in biological processes related to metabolism, transporters and angiogenesis in the COVID-TMA glomeruli. **C**) Dotplots of gene expression for selected genes. Each dot represents an individual glomerulus classified according to histology. For all comparisons of COVID-TMA to normal glomeruli, parametric ANOVA p< 0.05. Glomeruli from other patients/histologic profiles are included for completeness **D**) RNA *in situ* hybridization of VEGFA confirms its digital spatial expression profile shown in panel C.

## DISCUSSION

In this study, we successfully demonstrate feasibility of DSP to understand HIV and SARS-CoV-2 associated collapsing glomerulopathy and TMA. In contrast to the prior studies that only examined idiopathic collapsing glomerulopathy, we describe here the expression profile of collapsing glomerulopathy associated with viral infection.^8, 9^ We compared results using DSP to prior studies using laser capture microdissection and confirmed that collapsing glomerulopathy has a distinct expression profile with increased expression of developmentally regulated and ECM remodeling genes. However, the increased sensitivity and glomerulus-level resolution of DSP helped to identify differences in expression profiles when glomeruli were stratified by histology or disease status. For example, glomeruli from HIV-infected patients exhibited a strong interferon response signature that was not present in glomeruli from COVID-19 patients. This was surprising since we and others have proposed that COVID-19 associated collapsing glomerulopathy develops in the context of exuberant cytokine responses in individuals with high risk APOL1 genotypes.^5-7^ It is possible that this difference reflects the ongoing viral infections in 2/3 of the HIV patients, versus cleared infections in 2/3 of the COVID-19 patients. Similar to the reports of direct infection of kidney cells by HIV, establishment of direct SARS-CoV-2 infection of the kidney has proved controversial.^7^ We did not detect SARS-CoV-2 viral RNA in any of our samples by spatial profiling or RNA-ISH. The lack of evidence for SARS-CoV-2 infection in the biopsies supports an indirect injury model, but does not exclude the possibility of prior transient infection (“hit and run” model).^7^ We also identify a distinct vascular injury profile in TMA which has not been previous described. Collapsing glomeruli from HIV and COVID-19 patients exhibited almost complete loss of VEGFA which could represent downregulation of the gene and/or loss of podocytes. However, glomeruli from patient COVID3 with TMA showed increased expression of VEGFA and other endothelial cell related transcripts, which has not been previously described and could represent an adaptive response to vascular injury.

The main limitation of our study is the small number of samples; however, this is inherent to studying rare entities such as collapsing glomerulopathy. For comparison even Nephroseq, a large database with >2,100 patient datasets, only has expression data from 6 patients with collapsing glomerulopathy, the same number as in our study (www.nephroseq.org, accessed July 2021, University of Michigan, Ann Arbor, MI). Furthermore, the ability to correlate a single glomerulus’ histology to its expression profile is not technically possible with laser capture microdissection which is the most common method used in the Nephroseq dataset. Another shortcoming is the small number of genes available in the Cancer Transcriptome Atlas probeset that was available at the time we performed this study (1,825 cancer and immunology-focused genes). Since then, a Whole Transcriptome Atlas probeset encompassing ∼18,000 protein coding genes has been released which will enable whole transcriptome analyses of disease processes in kidney biopsies. Other limitations of our study include validation failures if glomeruli were not preserved on deeper levels used for RNA-ISH or immunohistochemistry and difficulty in establishing causality and timing of mechanisms, which is common in all human biopsy-based studies. However, even with these limitations, we uncovered remarkable heterogeneity in glomerular expression patterns by disease and histology using DSP. Importantly, we found that there was good correlation between DSP and RNA-ISH and protein levels determined by immunofluorescence/immunohistochemistry, validating this new technology and its utility for providing novel insight into the pathophysiology of disease in tissues.

Most powerfully, DSP technology is sensitive enough to perform studies on single sections of FFPE tissue with even small numbers of glomeruli and does not require pooling, thereby enabling single glomerulus level resolution data in any clinical or research biopsy. This is especially useful for rare disease processes or those with only focal involvement of kidney structures such as collapsing glomerulopathy. In these instances, the pooling of large numbers of structures to meet the minimum input requirements for laser capture microdissection-based profiling would obscure underlying expression patterns, as we have shown in this study. Another advantage of DSP’s probe-based mRNA quantification is compatibility with single sections of FFPE clinical archive specimens. By contrast, laser capture microdissection requires higher RNA input integrity and is preferentially performed on prospectively collected and stabilized samples from patients that have consented to have extra research biopsies performed specifically for these studies.

Spatially resolved profiling of gene expression represents the newest iteration of paradigm shifting technology that is poised to revolutionize our understanding of disease mechanisms.^33^ Emerging spatial transcriptomic methods^34, 35^ leverage widely available next generation sequencers to quantify transcripts from tissue sections applied to slides in simplified workflows. Recently, DSP has been applied to COVID-19 autopsy kidney tissues,^21^ but to the best of our knowledge, this study represents the first application of this technology to glomeruli in kidney biopsy tissues. In this study, DSP revealed glomerular heterogeneity and shared and distinct pathologic pathways in HIV and SARS-CoV-2 driven collapsing glomerulopathy and TMA. This opens the exciting possibility of using archival FFPE specimens to provide insight into a variety of anatomically defined kidney disease mechanisms using human biopsy tissues.

## FUNDING

This study was supported by a pilot research grant from the Department of Laboratory Medicine and Pathology at the University of Washington. S.A. also received support from NIH/NCATS 3 UG3/UH3TR002158-04S1 (PI Jonathan Himmelfarb).

## FINANCIAL DISCLOSURE

S.A. owns common stock of nanoString (NSTG), the manufacturer of the GeoMx digital spatial profiling instrument.

## SUPPLEMENTAL FIGURE & TABLES

**Supplemental Figure 1.**
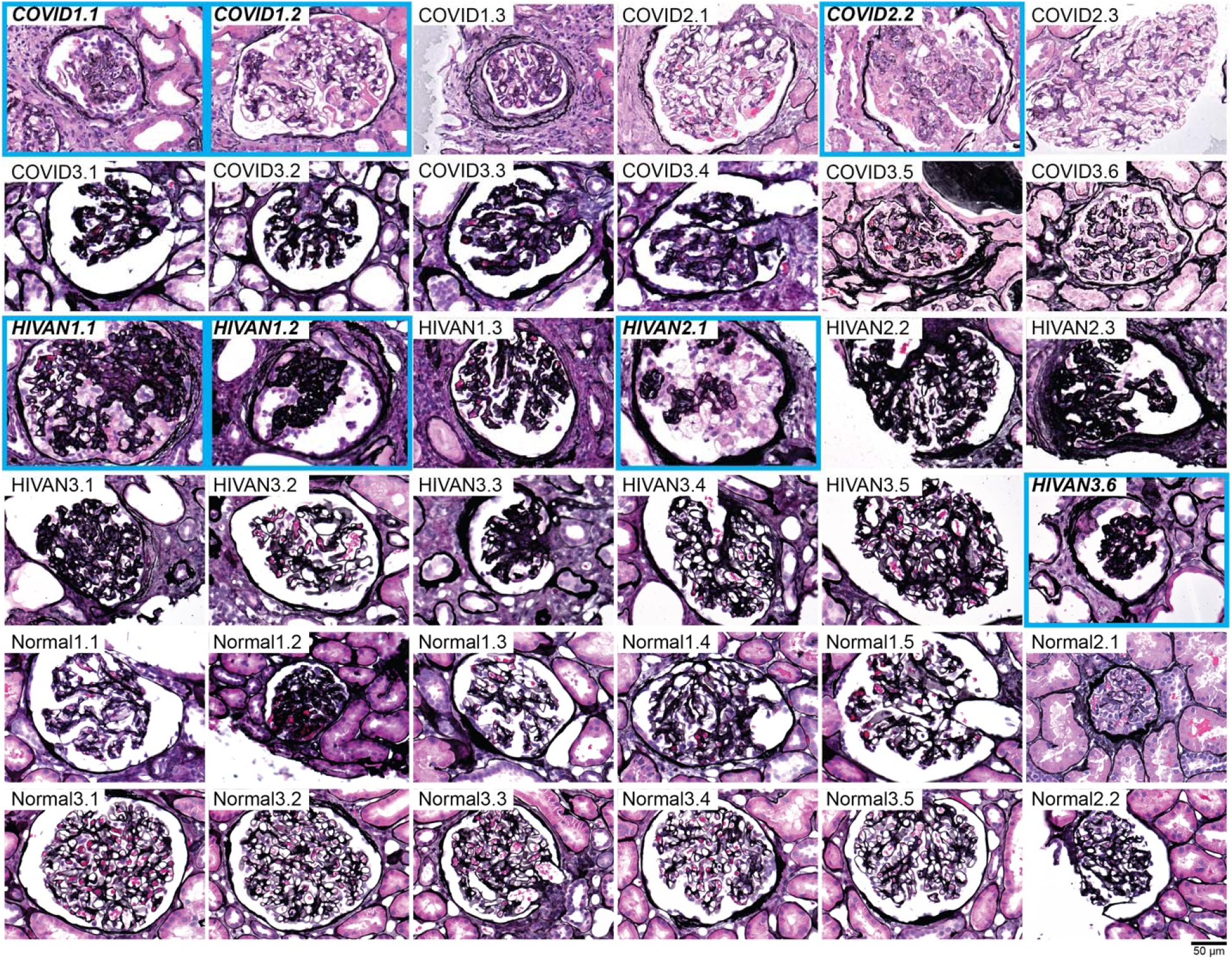
Histology of glomeruli selected for digital spatial profiling. For each glomerulus the first number indicates the patient and the second, the glomerular ID for that patient. Glomeruli with features of collapsing glomerulopathy are indicated in ***bold italics*** and a blue bounding box. Attempts were made to identify glomeruli on original 2µm thick Jones-stained sections. When this was not possible, the glomeruli were imaged from the slide used to generate spatial profiling data that was post-stained with Jones. Apparent differences in appearance/quality may be due to the fact that these sections were cut at 4µm and have been processed previously for digital spatial profiling.

**Supplemental Table 1.** Clinical and demographics of patients in this study.

| ID | Age | Sex | SCr (mg/dL) | Proteinuria | Clinical Presentation |
| --- | --- | --- | --- | --- | --- |
| Normal1 | 52 | F | 0.94 | No | gross hematuria, flank pain and negative urologic work-up |
| Normal2 | 48 | F | 0.60 | 1.3 (UPC) | recurrent gross hematuria |
| Normal3 | 28 | M | 0.70 | 2 (g/24hr) | gross hematuria, flank pain |
| HIVAN1 | 33 | M | 8.05 | 19.9 (UPC) | AIDS (non-compliant on HAART), CD4 count of 34, hypertension and CHF grade I, who developed nephrotic range proteinuria |
| HIVAN2 | 53 | M | 6.1 | 24.5 (UPC) | AIDS (Pneumocystis pneumonia >20 years ago) but now on HAART with undetectable viral load, presenting with kidney failure and proteinuria |
| HIVAN3 | 51 | M | 2.4 | 3+ (UA) | Recent diagnosis of HIV with low CD4 count, presenting with kidney injury |
| COVID1 | 47 | M | 6.6 (on dialysis) | 7.6 (UPC) | Acute kidney injury; At biopsy, COVID PCR negative 25 days after initial positive PCR |
| COVID2 | 44 | M | 12.0 | 11.4 (UPC) | Acute kidney injury; At biopsy, COVID PCR negative 6 weeks after initial positive PCR |
| COVID3 | 63 | F | 6 | Not checked | Oliguric acute kidney injury; At biopsy, patient was COVID PCR positive |

**Abbreviations:** AIDS – acquired immunodeficiency syndrome; CHF – congestive heart failure; HAART – highly active anti-retroviral therapy; SCr – serum creatinine; UA – urinalysis; UPC – urine protein to creatinine ratio.

**Supplemental Table 2.** Biopsy findings of patients in this study.

| ID | # glomeruli | # globally/segmentally sclerosed | IF/TA | Vascular findings | EM findings | Diagnosis |
| --- | --- | --- | --- | --- | --- | --- |
| Normal1 | 20 | 3 / 0 | Mild | None | No deposits, good podocyte foot process integrity | Minimal nephrosclerosis |
| Normal2 | 45 | 0 / 0 | None | None | No deposits, good podocyte foot process integrity | No significant histopathologic alterations |
| Normal3 | 31 | 1 / 0 | None | Mild arteriolar hyalinosis | No deposits, good podocyte foot process integrity | No significant histopathologic alterations |
| HIVAN1 | 20 | 15 / 4 | Moderate-to-severe; microcystic dilatation of atrophic tubules | Moderate arteriosclerosis | No deposits, diffuse podocyte foot process effacement | HIV-associated nephropathy |
| HIVAN2 | 16 | 10 / 4 | Severe | Moderate-to-severe arteriolar hyalinosis | No deposits, segmental podocyte foot process effacement | Collapsing FSGS |
| HIVAN3 | 26 | 11 / 5 | Moderate; microcystic dilatation of atrophic tubules | Moderate arteriosclerosis | No deposits, diffuse podocyte foot process effacement | HIV-associated nephropathy |
| COVID1 | 14 | 5 / 2 | Mild | Thrombotic microangiopathy | No deposits, segmental podocyte foot process effacement | Collapsing FSGS and thrombotic microangiopathy |
| COVID2 | 4 | 1 / 1 | Moderate | None by light microscopy | No deposits, segmental podocyte foot process effacement, subendothelial space widening | Collapsing FSGS and thrombotic microangiopathy |
| COVID3 | 36 | 0 / 3 | Minimal | Arteriolar and glomerular thrombi and endothelial cell swelling | No deposits, diffuse podocyte foot process effacement, subendothelial space widening and endothelial cell swelling | Collapsing FSGS and thrombotic microangiopathy |

Abbreviations: FSGS – focal and segmental glomerulosclerosis; IF/TA – interstitial fibrosis, tubular atrophy;

**Supplemental Table 3. Sequencing parameters for glomerular areas of interest (AOI) used in this study**. Sequencing parameters can be correlated to glomerular histology shown in Supplemental Figure 1.

**Supplemental Table 4. Q3 normalized gene expression for glomeruli in this study**. Gene expression can be correlated to glomerular histology shown in Supplemental Figure 1.

